# Community structure of known and previously unknown endobacteria associated with spores of arbuscular mycorrhizal fungi

**DOI:** 10.1101/2023.07.25.550273

**Authors:** Olga A. Lastovetsky, Tancredi Caruso, Fiona P. Brennan, David Wall, Susanna Pylni, Evelyn Doyle

## Abstract

Arbuscular mycorrhizal fungi (AMF) are ubiquitous plant root symbionts which can house two endobacteria: *Ca.* Moeniiplasma glomeromycotorum (*Ca*Mg) and *Ca*. Glomeribacter gigasporarum (*Ca*Gg). However, little is known about their distribution and population structure in natural AMF populations and whether AMF can harbour other endobacteria. We isolated AMF from two environments and surveyed the surface-sterilized spores for endobacteria. We found that *Ca*Mg and *Ca*Gg differed significantly in distribution whereby *Ca*Mg were extremely abundant (80%) and *Ca*Gg were extremely rare (2%) in both environments. Unexpectedly, we discovered an additional and previously unknown level of bacterial diversity within AMF spores which extended beyond the known endosymbionts, with as many as 277 other bacterial taxa detected in individual spores. Detailed analysis of endobacterial communities inside AMF spores revealed that: (i)*Ca*Gg were not limited in distribution to the *Gigasporacea* family of AMF, as previously thought, (ii) *Ca*Mg community structure was driven by AMF host genotype, (iii) a significant inverse correlation existed between the diversity of *Ca*Mg and diversity of all other endobacteria. The latter suggests the existence of competition dynamics between different bacterial populations inside AMF spores and provides a basis for generation of testable hypotheses regarding the function of *Ca*Mg in AMF biology.

## Introduction

The majority of land plants, including major crop species, form mutualistic symbiotic associations with arbuscular mycorrhizal fungi (AMF, Glomeromycotina)^1^. AMF associate with plant roots, providing their host with important mineral nutrients in exchange for carbon^1^. Through this nutrient exchange as well as their roles in disease protection, water uptake and soil aggregation, AMF are important components of ecosystem functioning and are of major interest in sustainable agriculture^2 3–5^.

AMF themselves are known to harbor intracellular bacteria (i.e. “endobacteria”) which live within fungal hyphae and spores^6^. Two different AMF endobacteria have been described to date, *Ca.* Moeniiplasma glomeromycotorum (*Ca*Mg, Tenericutes)^7^ and *Ca*. Glomeribacter gigasporarum (*Ca*Gg, β-proteobacteria)^8^. These endobacteria are vertically-transmitted in fungal spores and differ significantly in population structure, distribution and molecular evolution patterns.

*Ca*Gg are the best-studied endobacteria and are believed to be conditional mutualists of AMF. *Ca*Gg prime the energy metabolism of AMF^9^ and increase hyphal proliferation during spore germination^10^ as well as sporulation success of their host^9^. The evolutionary history of *Ca*Gg indicates significant patterns of co-evolution with AMF^11^. *Ca*Gg populations are homogenous within AMF individuals^11^ meaning that only a single *Ca*Gg haplotype is found within an AMF isolate. Despite their benefit to AMF,*Ca*Gg appear rare in nature^12^ and have only been found in AMF belonging to the family *Gigasporacea*^13^.

*Ca*Mg endobacteria are more common in AMF, being found in association with phylogenetically diverse AMF^14^ and in a high proportion of environmental AMF isolates^12^. Like *Ca*Gg, *Ca*Mg are ancient associates of AMF^15^ but their role in fungal biology remains uncharacterized due to their uncultivability and difficulty in obtaining “cured” isolates. However, their molecular evolution patterns and metabolic profile indicate that*Ca*Mg might be antagonists of AMF^16, 17^. *Ca*Mg populations within AMF isolates are genetically-diverse^7, 12, 14, 16–18^ *i.e.* a single AMF spore can harbour multiple *Ca*Mg haplotypes. This intra-host diversity is thought to be maintained by various genetic recombination machinery encoded in their genomes^15^ as well as occasional horizontal transmission^16^. These molecular evolution characteristics are unusual for vertically-transmitted mutualists, as high intra-host diversity is expected to increase symbiont competition and exploitation of host resources, as well as increase rates of horizontal transmission to secure new hosts^19^. Moreover, *Ca*Mg are closely related to bacteria with parasitic lifestyles including *Mycoplasma pneumoniae*, a parasite of animals^17^ and Mycoplasma-related endobacteria of Mortierellamycotina fungi which are antagonistic to their fungal host under certain laboratory conditions^20^. Despite these characteristics of parasites, it remains unresolved why *Ca*Mg are maintained within AMF populations to such high levels if they do not provide any benefit to AMF, as it is expected for selection to remove such parasites from populations^21^.

Most of what we know about AMF-associated endobacteria comes from studies of culture collection AMF isolates with only one study to date focusing on this relationship in nature^12^. Moreover, previous studies have largely focused on known AMF endobacteria using *Ca*Gg- and *Ca*Mg-specific primers, thus overlooking any other potential bacterial associates. In this study, we aimed to (1) characterize the entire diversity of endobacteria associated with environmental AMF isolates; (2) determine which factors (e.g. soil chemistry, host genotype) affected the distribution of endobacteria in AMF and (3) conduct and in-depth population structure analysis of *Ca*Mg endobacteria to better understand their role in AMF biology. We isolated AMF spores from two sampling sites: a natural dune and an agricultural grassland located in Co. Wexford, Ireland. We surveyed the isolated AMF spores for presence of known endobacteria and used modelling to determine which environmental parameters affected their distribution in AMF. Next, using deep sequencing, we characterized the entire diversity of bacteria found within AMF spores and analyzed their population structure with a focus on *Ca*Mg.

## Materials & Methods

### Sampling site description and strategy

Sampling was conducted at two sites located within 13 km of each other on the east coast of Ireland. Site 1 was an agricultural grassland located at Teagasc, Johnstown Castle, Co. Wexford, Ireland (52° 17’ 55’’ N, 6° 29’ 49’’W). This site was seeded with *Lolium perenne* (perennial ryegrass), divided into 16 plots in a randomized block design and subjected to different levels of long-term phosphorus (P) fertilization (since 1995) and slurry amendments (since 2016) (Figure S1). Each P treatment received annual fertilisation rates of 0, 15, 30 and 45 kg ha^-1^ yr^-1^ of 16% superphosphate (2 CaSO_4_ + Ca(H_2_PO_4_)_2_), hereafter referred to as P0, P15, P30 and P45. The aboveground plant material was harvested eight times per year to simulate grazing. Soil samples were collected in May and November 2018 from plots untreated with slurry (“-slurry” plots, Figure S1) using a soil corer (AMS 404.45 soil core sampler and slide hammer, American Falls, ID, USA). Three soil cores were obtained from each plot and combined into a single composite sample. Additionally, small cores (using AMS 56975 soil probe, American Falls, ID, USA) were collected from each plot and combined into a single composite sample for soil chemistry analysis.

Site 2 a dune located at Curracloe, Co. Wexford, Ireland (52° 23’ 20’’ N, 6° 21’ 45’’W). Samples were collected in May, August and November 2018. During May and November sampling four transects were laid out starting from the edge of vegetation at the seaward side (corresponding to the end of the beach) and extending 100 m inland (Figure S1).

Samples were taken every 20 m by collecting the soil around a single *Ammophila arenaria* (marram grass) plant. Samples were not collected from the 0 m mark (closest to the beach) as a result of sand burial and dry soil conditions. At each sampling point we recorded the distance to four nearest plants in the 70 cm radius around, later averaged to give average nearest neighbour distance (NND). Average NND was used as a measure of plant density. Sampling in August was performed along one 100 m long transect (n=5 samples) for AMF spore extractions in order to gauge peak spore abundance.

All samples were transported on ice to University College Dublin, Ireland. Soil was dried at room temperature and subsequently stored at 4°C before AMF spore extraction.

### Soil chemistry analysis

Soil chemistry analysis was performed at the Teagasc Soil laboratory at Johnstown Castle, Wexford, Ireland. Detailed methods are described in Lastovetsky *et al.* ^22^. In brief, soil pH was determined with a pH probe (WTW, Germany) using a 2:1 deionised water to soil mixture. The Mehlich III method ^23^ was used to analyse soil nutrient availability (P, K, Al, Ca, Co, Cu, Fe, Mg, Mn, S, Zn and Cu).

### Spore extraction, collection and decontamination

AMF spores were extracted from 50 g of air-dried soil suspended in 200 ml water using wet sieving and decanting followed by 2M sucrose centrifugation as described by Daniels and Skipper ^24^. Spores were collected on 0.45 μm gridded nitrate cellulose filters (Whatman) and picked using fine tweezers under a dissecting microscope. Due to low number of viable spores, all spores that appeared healthy were picked for further analysis. Selected spores were decontaminated individually as in Mondo *et al.* ^11^. In brief, spores were subject to sequential washes with 1 mM and 50 mM H_2_O_2_, then with 4% chloramine T, followed by two final washes with sterile nanopure water.

### AMF spore Identification

Following surface decontamination, each spore was crushed with a pipette tip to release its contents. Total spore DNA was whole-genome amplified using Illustra^TM^ GenomiPhi-V2 kit (GE Healthcare), and the 1/20 diluted product was used for PCR. A fragment of the fungal 18S ribosomal RNA (rRNA) gene was PCR amplified from individual spores with primers AML2^25^ (GAACCCAAACACTTTGGTTTCC) and WANDA^26^ (CAGCCGCGGTAATTCCAGCT) using JumpStart RedTaq DNA Polymerase Master Mix (Sigma). PCR products were sent for cleaning and Sanger sequencing (Macrogen Europe). DNA sequences were edited in Geneious Prime V11 (Biomatters Ltd), aligned with MUSCLE^27^, and grouped into operational taxonomic units (OTUs) at 95% similarity cutoff using mothur^28^. To assign AMF identity to each spore, we performed phylogenetic analyses to cluster the representative sequences from each OTU with reference AMF sequences into statistically supported virtual taxonomic units (VTUs) as described previously^12^.

### Screening for endobacteria

AMF spores were screened individually for incidence of *Ca*Gg by PCR with *Ca*Gg/Burkholderia-specific primers GlomGiGF^13^ (GGGTCCATTGCGGATTACTTC) and LSU483r^11^ (GGTGCAGGAATATTAACC) amplifying a portion of the 23S rRNA gene, followed by Sanger sequencing, as described above. *Ca*Mg were detected by gel electrophoresis of PCR products generated with *Ca*Mg-specific primers 109F-1 (ACGGGTGAGTAATRCTTATCT), 109F-2 (ACGAGTGAGTAATGCTTATCT), 1184R-1 (GACGACCAGACGTCATCCTY), 1184R-2 (GACGACCAAACTTGATCCTC), 1184R-3 (GATGATCAGACGTCATCCTC)^14^ targeting a portion of the 16S rRNA gene. Spores were also screened for other endobacteria using PCR the universal bacterial primers 27F (AGAGTTTGATCMTGGCTCAG) and 1492r (TACGGUTACCTTGTTACGACT) amplifying a portion of the 16S rRNA gene, followed by gel electrophoresis.

### Statistical analysis

Linear regression was used to examine the relationship between different soil chemistry characteristics (soil nutrients and pH, data presented in Table S1) in R (v. 4.0.3) with the*lm* and *abline* functions. Influence of environmental parameters on *Ca*Mg incidence in AMF spores was analyzed using general linearized mixed models with binomial distribution with the *lsmeans* and *lme4* packages in R. Environmental variables (plant density, soil nutrient content, pH) were modelled as fixed effects. To account for variability between and within plot blocks (Site 1, see Figure S1) and transects (Site 2), these were modelled as random effects. All environmental parameters were modelled as continuous variables.

### MiSeq Illumina library preparation and sequencing

DNA from spores which tested positive for presence of any kind of bacteria in our PCR screen (n=87) were prepared for Illumina Miseq sequencing using dual index primers in a one-step PCR method as described in Kozich et al^29^. Briefly, the V4 region of the bacterial 16S ribosomal RNA (rRNA) gene was PCR amplified with primers 515F (GTGCCAGCMGCCGCGGTAA) and 806R (GGACTACHVGGGTWTCTAAT) containing Illumina adapter sequences and index barcodes. Each 20 µl PCR reaction contained 10 µl of JumpStart RedTaq DNA Polymerase Master Mix (Sigma), 8 µl dH_2_O, 1µl of indexed primers (0.05 µM final concentration) and 1 µl template DNA. PCR conditions were as follows: 94°C for 2 mins followed by 35 cycles of 94°C for 30s, 55°C for 30s, 72 for 2 mins followed by a final extension at 72°C for 5 mins. All PCR reactions were conducted in duplicate and subsequently pooled. PCR products were cleaned and normalized using the SequalPrep Normalization Plate Kit (Invitrogen, Massachusetts, USA) according to manufacturer’s instructions. The amplicon pool was sent to the Centre for Genomic Research, University of Liverpool for sequencing on the Illumina Miseq 250 PE platform.

### Bioinformatic analysis

#### Sequence processing

The raw Fastq files were trimmed for the presence of Illumina adapter sequences using Cutadapt version 1.2 ^30^. The option -O 3 was used, so the 3’ end of any reads which match the adapter sequence for 3 bp or more were trimmed. Reads were further trimmed using Sickle version 1.2 (https://github.com/najoshi/sickle/releases/tag/v1.2) with a minimum window quality score of 20. Reads shorter than 15 bp after trimming were removed. If only one of the read pairs passed this filter, both reads were removed from downstream analysis.

Trimmed reads which passed quality control were processed following the mothur Miseq standard operating procedure, SOP^29^. Briefly, each read was trimmed to a maximum of 330 bp, ambiguous bases were removed and sequences containing homopolymer runs >8 bases were discarded. Reads were aligned to the SSU SILVA reference alignment v132 customized to the 16S V4 region followed by identification and removal of chimeric sequences using the VSEARCH algorithm. 16S sequences were classified using the classify.seqs command and the mothur-formatted RDP Classifier reference trainset v.18 files, customized to include known AMF endobacterial sequences. 16S sequences were then clustered into operational taxonomic units (OTUs) at 0.03 cutoff using the ‘cluster’ command with default opticlust method. For analysis of *Ca*Mg population structure, sequences belonging to the Tenericutes phylum were re-clustered into OTUs at 0.06 cut-off to represent 94% 16S similarity level as is the standard for this group of bacteria^7^.

On average 17 453 reads were obtained from each sample. The data was further rarefied to an even depth of 6 477 corresponding to reads in a sample with lowest number of reads. It was not possible to identify *Ca*Gg sequences in our dataset following the mothur MiSeq SOP (classify.seqs command using the default wang method) even after customizing the taxonomy and reference files to include known *Ca*Mg and *Ca*Gg sequences. The classify.seqs command identified only a small subset of *Ca*Gg-related sequences. We therefore took a manual approach and analysed all sequences assigned to “betaproteobacteria” in the spores which were positive for *Ca*Gg in our initial PCR screen with *Burkholderia*-specific primers (samples 705_P30_2 and T2_100_13). These spores harboured two OTUs with hits to *Ca*Gg in the NCBI BLASTn database (OTU0094 and OTU0138). Using these OTUs as references, we queried the NJ tree of all the “betaproteobacteria” OTUs using the phyloseq plot_tree function. This identified a further two putative *Ca*Gg OTUs (OTU0077 and OTU0034) which grouped close to OTU0094 and OTU0138 and an additional 12 OTUs which grouped further away. Using BLASTn we pulled out the OTUs which showed sequence similarity to *Ca*Gg and/or known closely related fungal endobacteria *Mycoavidus cysteinexigens* and *Mycetohabitans rhizoxinica* (Figure S2). Using Bayesian and Maximum Likelihood phylogeny reconstruction (Figure S3) we confirmed their close affiliation to *Ca*Gg giving a total of 5 *Ca*Gg OTUs (Otu0034, Otu0077, Otu0094, Otu0116, Otu0138) recovered from 11 different AMF spores.

#### Multivariate Analysis

Bacterial community analyses were conducted in R v. 4.0.3 with the phyloseq and vegan package^31, 32^. Separation of bacterial communities by location/AMF genotype was visualized using Principle Coordinate Analyses, PCoA and the significance of separation was tested based on Bray-Curtis dissimilarity distances using a permutational analysis of variance, PERMANOVA (adonis function). Differences in microbial diversity (Shannon diversity index) were conducted using a pairwise Wilcoxon rank sum test through the pairwise.wilcox.test function.

### Phylogeny reconstructions

For the generation of the AMF, *Ca*Gg and “unclassified bacteria” phylogenies, DNA sequences were aligned using MUSCLE^27^. Phylogenies were reconstructed under the GTR+I+Γ nucleotide substitution model implemented in MrBayes 3.2.7^33^, with a 25% burn-in and the average standard deviation of split frequencies (<0.01) used as a convergence diagnostic. Maximum likelihood phylogenies were created with PhyML 3.3.2 implemented from within Geneious Prime V11 (Biomatters Ltd) under the GTR+I+Γ nucleotide substitution model with 1000 bootstraps.

### Phylogenetic diversity estimates

For the generation of phylogenetic distance estimates of *Ca*Mg communities inside AMF spores, the representative sequences from all *Ca*Mg OTUs were aligned using MAFFT^34^. Phylogeny was reconstructed using FastTree^35^ with the GTR + CAT model of nucleotide substitution. We quantified the phylogenetic structure^36^ of the community inside the AMF by crossing the information in the phylogenetic tree with that of the OTU by sample matrices. We calculated the Faith phylogenetic diversity metrics^37^and the mean pairwise and nearest taxon distance^38^ using the R picante package^39^ With the same package we also calculated the effect size of the deviation of the observed value of the metric from a null model. We used the ‘independentswap’ with 999 replicates, as this algorithm retains column and row totals as observed in the species co-occurrence, which is best suited to test for purely compositional effects while keeping features such as richness in the observed matrix^40^.

## Results

### AMF spore-based community structure

Spore-based studies of AMF are lacking in Ireland therefore we first set out to identify the optimum sampling time for peak spore abundance. In the American North Atlantic dunes, spores peak in abundance in November-December^41, 42^, but in Ireland it was expected that peak spore abundance could be anytime between May and November^43, 44^. Spores isolated in May 2018 were mostly degraded and parasitized: out of ca. 60 spores that were processed, only 6 were successful in amplifying AMF. August 2018 sampling revealed presence of only degraded and parasitized spores, whereas November 2018 sampling identified a high number of viable spores. These data indicate that analogous to what has been previously observed in the American N. Atlantic dunes, AMF spore abundance in Irish dunes peaks in late autumn-early winter^12, 45^. It is important to note, however, that because we did not conduct any sampling between November and May, AMF spore abundance might be even higher during that time.

In total, we extracted and processed 467 spores, 240 from site 1 (agricultural grassland) and 227 from the site 2 (dune). The success rate of genotyping spores was 15% from the agricultural grassland and 26% from the dunes. Spores were grouped into 9 virtual taxonomic units (VTUs) at the genus level using a combination of sequence similarity and Bayesian as well as maximum likelihood phylogeny reconstruction following a previously established method^12^ (Figure 1). Agricultural grassland spore AMF communities were composed of 5 VTUs and were dominated by VTU *Funneliformis*, the dune communities were composed of 4 VTUs dominated by VTU *Diversispora* (Figure S4).

**Figure 1.**
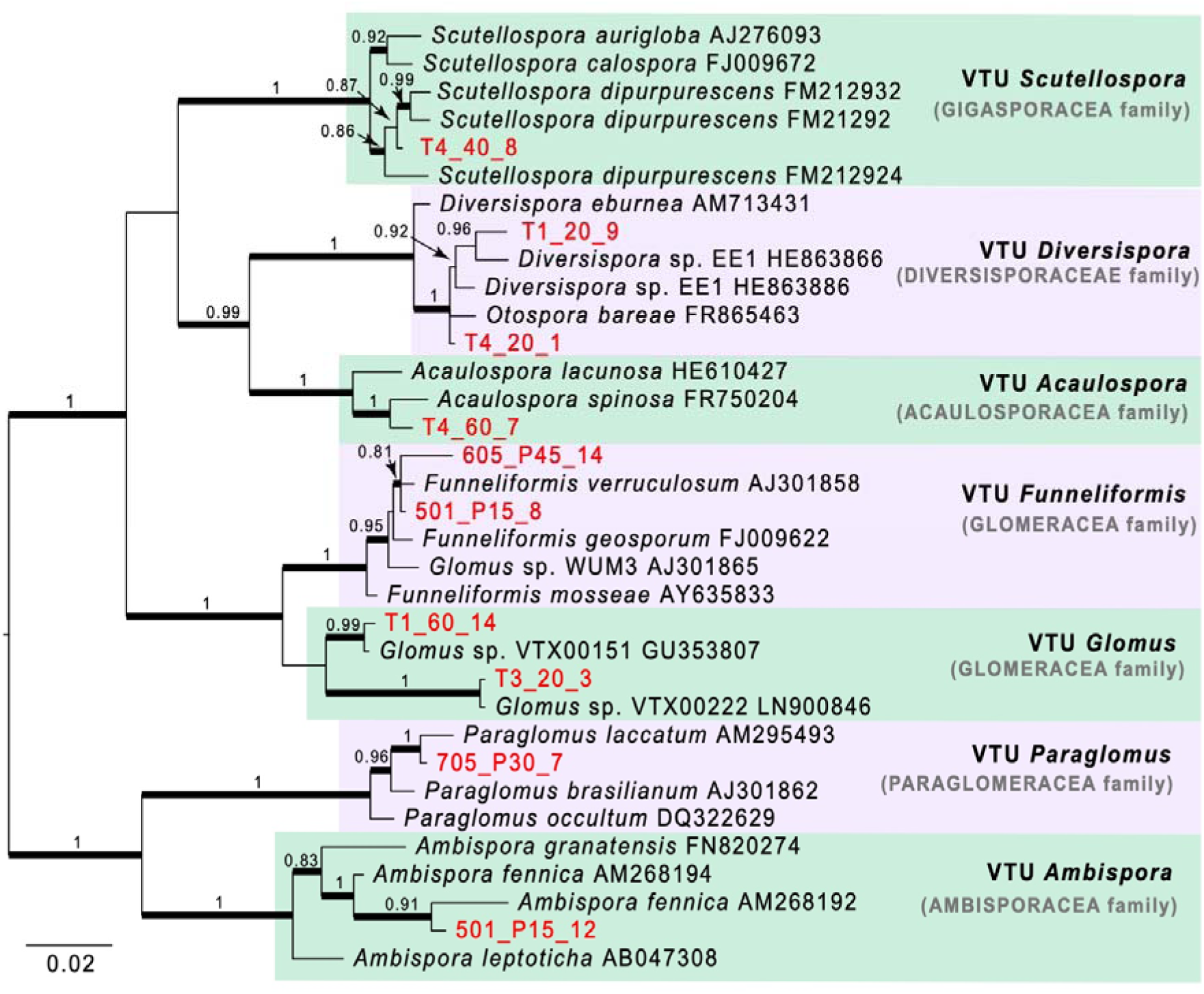
AMF phylogeny based on the 18S rRNA gene. Taxa in red are representative OTUs from this study clustered at 97% sequence similarity, the remaining are reference taxa. Taxa identifiers include transect/plot number, sample number/P application rate and spore number. Bayesian posterior probability values >0.8 are displayed above branches, branches with Maximum Likelihood bootstrap support >70% are thickened. The tree was midpoint rooted.

### Effect of environmental parameters on distribution of known endobacteria across AMF spores

We screened all spores, which were surface sterilized prior to molecular work, for presence of known endobacteria using PCR with *Ca*Gg- and *Ca*Mg-specific primers. *Ca*Mg were detected in the majority of spores isolated from dunes (78%) and the agricultural grassland (80%). *Ca*Gg on the other hand, were detected only in one spore each from the two sites. Additionally, our PCR screen identified 5 spores which did not amplify*Ca*Gg or *Ca*Mg, but amplified a band with universal bacterial primers, indicating that these spores harboured previously uncharacterized endobacteria.

*Ca*Mg distribution differed among the AMF VTUs (Figure S5). *Ca*Mg was highly abundant (>80%) in the majority of spores (VTU *Diversispora* 100%, VTU *Paraglomus* 100%, VTU *Ambispora* 100%, VTU *Glomus* 100%, VTU *Scutellospora* 86% and VTU *Funneliformis* 79%), except in spores of VTU *Acaulospora* where it was found in less than half of the spores (47%).

In site 1, distribution of *Ca*Mg was significantly affected by soil nutrient levels. For example, probability of harboring *Ca*Mg increased with increasing available soil P levels (*P*=0.011) (Figure 2). Probability of harbouring *Ca*Mg was also significantly affected by Ca (*P*=0.032) and S (*P*=0.01) levels as well as marginally affected by K (*P*=0.058), Fe (*P*=0.062) and soil pH (*P*=0.06). Some of these soil nutrients were in turn significantly correlated with P and pH (Figure S6). It is thus apparent that soil nutrient levels and pH affected*Ca*Mg distribution in AMF, but it was not possible to pinpoint which was the main driver since they themselves were correlated to one another. In turn, AMF spore communities differed slightly between low (P0, P15) and high (P30, P45) P treatments (Figure S7). Therefore, it is possible that the effect of soil nutrient levels on *Ca*Mg distribution is related to the effect of soil parameters on the structure of the AMF communities themselves.

**Figure 2.**
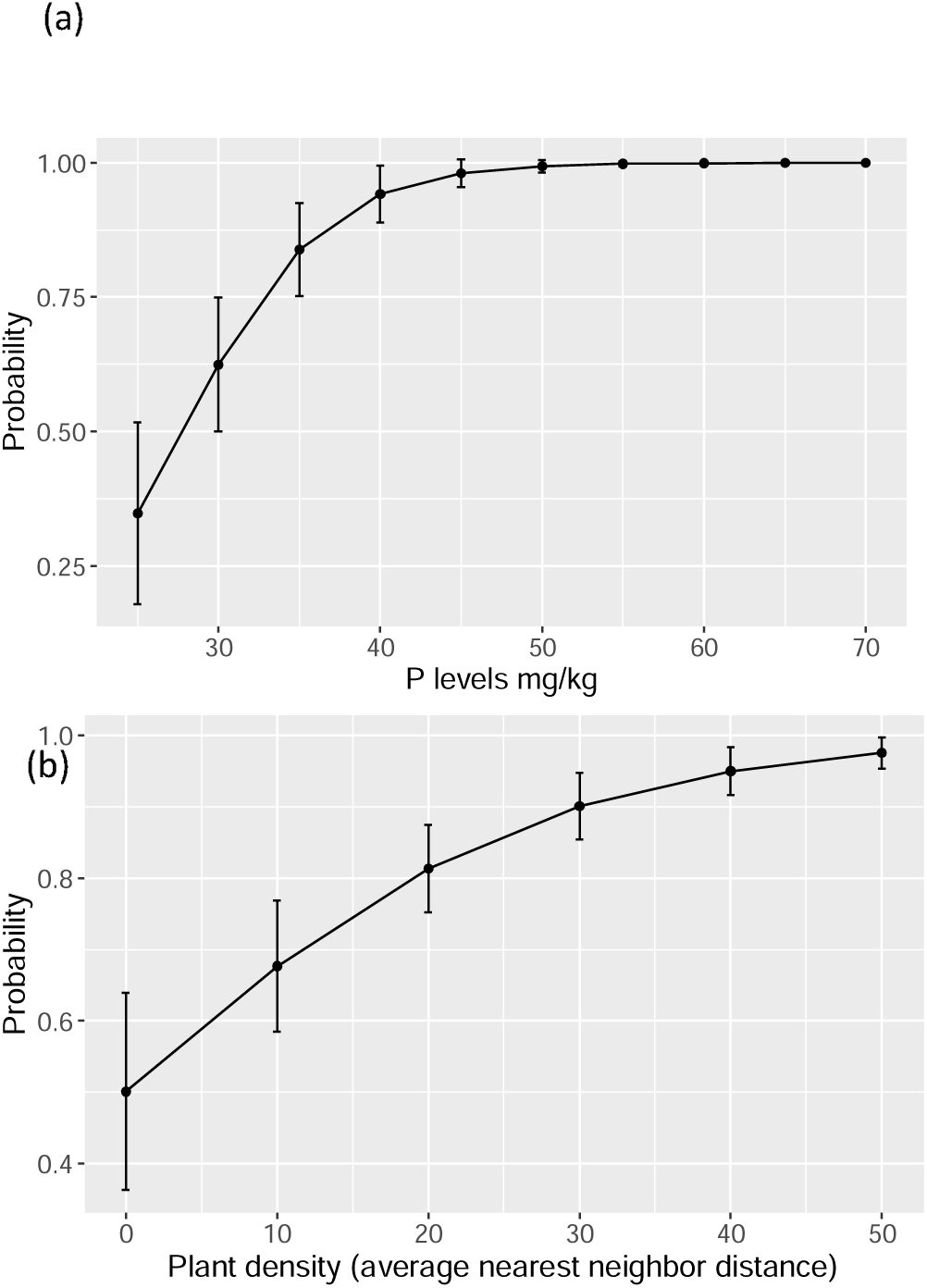
Distribution of *Ca*Mg in AMF spores at the agricultural grassland site. (a) Probability of AMF spores harbouring *Ca*Mg at different levels of soil P. (b) Probability of AMF spores harbouring *Ca*Mg at different plant density. Error bars represent 1 SEM.

Soil nutrient levels did not affect *Ca*Mg incidence in site 2, however plant density played a significant role (*P=*0.002) with higher probability of *Ca*Mg in areas of lower plant density (Figure 2).

Overall, it is apparent that multiple factors including soil nutrient levels, plant density and host genotype play a role in driving the distribution of *Ca*Mg endobacteria in AMF spore populations.

### Diversity and community structure of endobacteria within AMF spores

In order to understand the entire diversity of endobacteria associated with AMF spores, we characterized the bacterial communities from 84 spores (agricultural grassland: n=30, dunes: n=54; May: n=6, November: n=78). Rarefaction curves showed a surprisingly high level of diversity associated with each spore (Figure S8) with an average of 148 OTUs detected within individual spores. Diversity did not differ between the two sampling sites (Figure 3) but endobacterial community structure showed significant separation by site (PERMANOVA, *P*<0.001) and by host genotype (PERMANOVA, *P*<0.001) (Figure 3).

**Figure 3.**
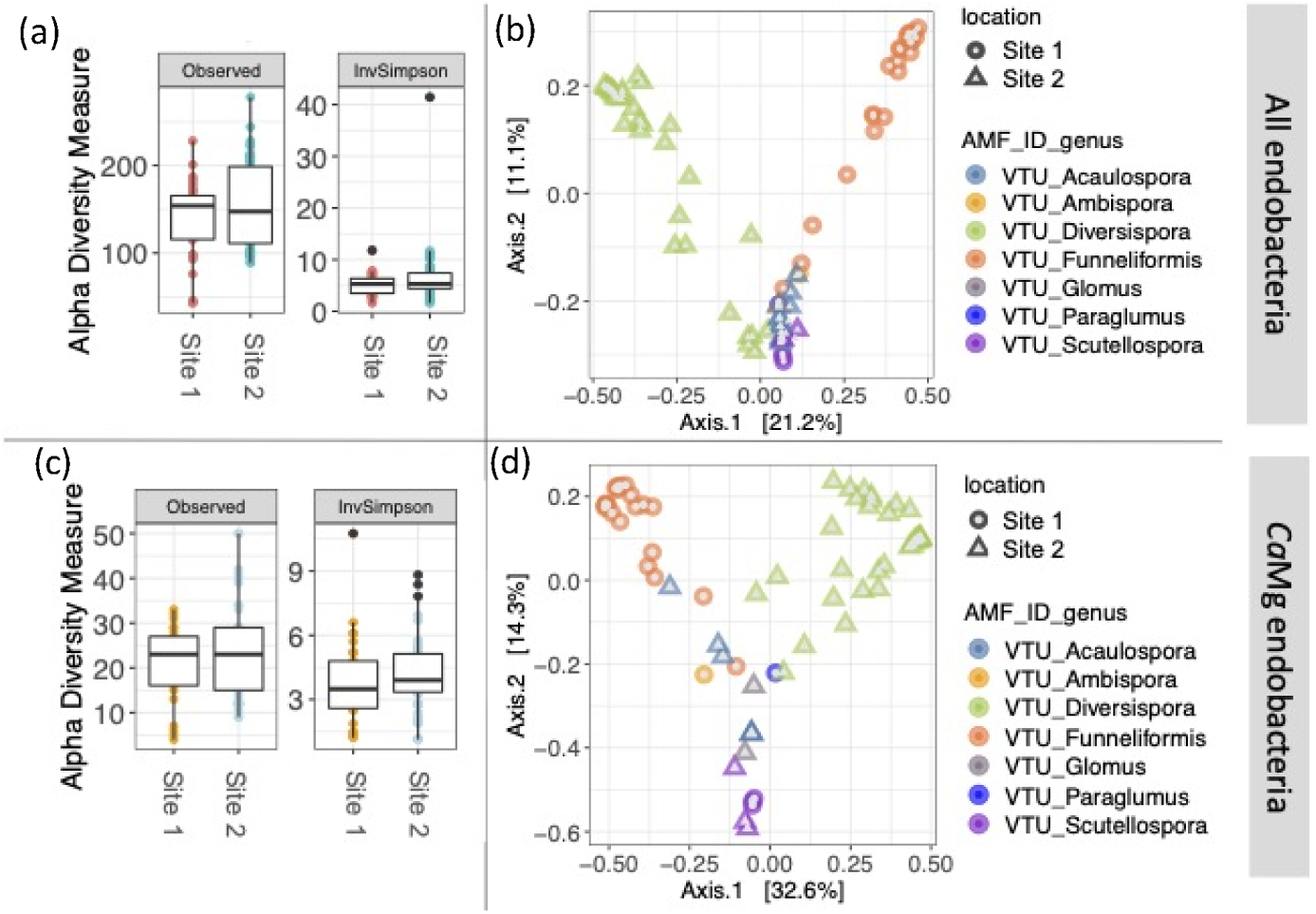
Diversity of bacteria inside AMF spores. (a) Total endobacterial richness (observed) and inverse Simpson (InvSimpson) measures of diversity in AMF spores; (b) Principal Coordinate Analysis (PCoA) plot of endobacterial communities inside AMF spores. PCoA was performed on a Bray-Curtis distance matrix. Each point represents an endobacterial community found within a single spore; (c) *Ca*Mg endobacterial richness and inverse Simpson measures of diversity in AMF spores; (d) Principal Coordinate Analysis (PCoA) plot of *Ca*Mg endobacterial communities inside AMF spores. PCoA was performed on a Bray-Curtis distance matrix. Each point represents an endobacterial community found within a single spore. Site 1, agricultural grassland; site 2, dune.

*Ca*Mg were extremely abundant within individual AMF spores, averaging to 82% relative abundance and representing >90% of reads associated with 55 out of 84 spores. Using deep sequencing, *Ca*Gg were detected in 10 spores, albeit only 6 spores had >500 *Ca*Gg reads. *Ca*Gg relative abundance within spores was 9% on average ranging from 0-27% of total reads (Table S2). These results are in agreement with previous qPCR and microscopy-based observations whereby *Ca*Mg were more abundant inside AMF spores than *Ca*Gg^6^. Notably, *Ca*Gg were only detected in spores with *Ca*Mg. Overall, *Ca*Mg were the most common and most abundant endosymbionts found inside AMF spores, whereas*Ca*Gg were extremely rare. In fact, it would have been impossible to detect *Ca*Gg in our dataset if it was not already a well-characterized associate of AMF.

Relative abundance of *Ca*Mg within AMF spores was related to host genotype and mirrored the overall distribution of *Ca*Mg across spores. *Ca*Mg showed the lowest relative abundance in VTU *Acaulospora* (37% on average) (Figure S9) which was also the VTU with lowest incidence of *Ca*Mg across all spores (47% of VTU *Acaulospora* spores harboured *Ca*Mg). Spores belonging to VTU *Glomus* also had comparatively low abundance of *Ca*Mg, whereas spores belonging to VTUs *Ambispora*, *Diversispora*, *Funneliformi*s and *Scutellospora* had >80% *Ca*Mg. Taken together these results indicate that host genotype exerts a level of control over *Ca*Mg endosymbiont populations having an effect on both the incidence and relative abundance of *Ca*Mg within spores.

Taking a closer look at the population structure of *Ca*Gg, we identified five different *Ca*Gg-related OTUs in our dataset. Three OTUs (OTU0138, OTU0034, OTU0094) clustered with known *Ca*Gg sequences, whereas the other two OTUs were more closely related to endosymbionts of *Mortierella elongata* (OTU0077) and *Rhizopus microsporus* (OTU0116) (Figure S3). As expected, only one *Ca*Gg genotype was found within individual AMF spores, confirming that this endosymbiont exists as a homogeneous population within AMF isolates ^11, 12^. To date, *Ca*Gg has only been detected in AMF belonging to the *Gigasporacea* family^11^. In our study, in addition to identifying *Ca*Gg reads in VTU *Scutellospora* and VTU *Diversispora* (both belong to the Gigasporacea family) we also detected *Ca*Gg in spores of VTU *Paraglomus* and VTU *Ambispora* (Table S2). To our knowledge, this is the first report of *Ca*Gg outside of Gigasporacea.

Interestingly, *Ca*Mg and *Ca*Gg were by far not the only endosymbionts found in AMF spores, with endobacteria belonging to 10 other Phyla detected in our dataset (Figure 4). Individual AMF spores were home to as many as 277 bacterial OTUs (mean=25) which were not *Ca*Mg or *Ca*Gg. Six spores were *Ca*Gg- and *Ca*Mg-free (<3% of *Ca*Mg reads, no *Ca*Gg reads), and these spores were instead dominated by endobacteria belonging to Acidobacteria, Proteobacteria and Verrucomicrobia (Figure S10). There were also spores in which *Ca*Mg dominated the endobacteria population, but which also contained reads from other bacterial taxa. Of the OTUs that did not belong to *Ca*Mg or *Ca*Gg, OTU0035 (Gammaproteobacteria, closest BLAST hit: *Escherichia coli* 214-4) was most commonly found in AMF spores (Figure S11). It was present in 76 out of 84 spores, with 10 spores having >100 OTU0035 reads. Other bacteria detected in AMF spores included members of Verrucomicrobia (with closest BLAST hits to uncultured soil bacteria), Alphaproteobacteria (*Rhizobeales*, *Rhodospiralles*) Gammaproteobateria (*Escherechia*, *Aquicella*, *Pseudomonas*) Actinobacteria and Acidobacteria. These bacteria are mostly known as free-living in terrestrial and aquatic environments, with some of them known to be facultative symbionts of plants (e.g. *Rhizobiales*) or parasites of animals (e.g. *Escherichia*) and protozoa (e.g. *Aquicella*).

**Figure 4.**
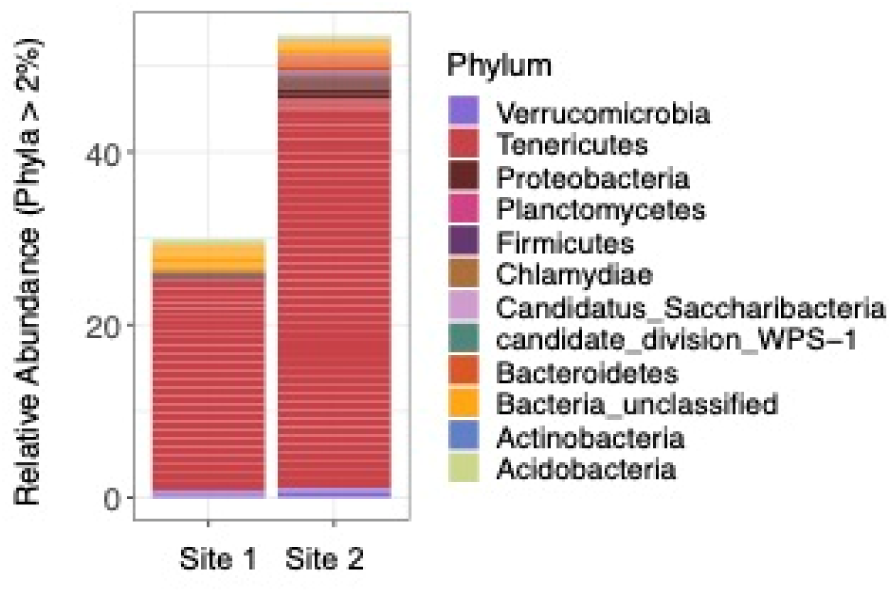
Relative abundance of endobacteria at Phylum level in AMF spores. The bars are of uneven heights due to less spores recovered from site 1. Site 1, agricultural grassland; site 2, dune.

In addition, there were a number of bacterial OTUs which were not classified at the Phylum level, and instead were only identified as belonging to “Bacteria”. These included highly abundant OTUs such as OTU0003 and OTU0066 with a total of 25 unclassified OTUs with >100 reads. We attempted to classify these using Bayesian and maximum likelihood phylogeny reconstruction and found a previously undescribed group of bacteria belonging to Mollicutes, which grouped together with *Mycoplasma*, *Spiroplasma* and *Ca*Mg but represented its own distinct clade (Figure S12). This new group was represented by 10 OTUs including the highly abundant OTU003 and OTU0066. The remaining 18 unclassified OTUs had hits only to sequences from environmental samples, and therefore it was not possible to conclusively determine which phylum they belonged to.

### Population structure of *Ca*Mg endobacteria

To better understand the diversity of *Ca*Mg within AMF spores, we re-clustered the reads associated with *Ca*Mg (all reads classified into the Tenericutes phylum) into OTUs at 94% sequence similarity, as is the standard for this taxonomic group^46^. We found that as many as 50 *Ca*Mg OTUs could be found in a single AMF spore, with an average of 22 OTUs per spore (Figure 3). This is in line with what is known about the population structure of these endosymbionts whereby multiple divergent genotypes can co-exist within individual AMF spores^7, 20^. The high level of diversity, however, was unprecedented because in the past no more than 3 OTUs clustered at 94% sequence similarity (or 15 haplotypes) were detected in individual AMF spores^16^.

*Ca*Mg diversity did not differ between spores isolated from different sites (Figure 3). However, *Ca*Mg richness did differ among the four main AMF VTUs (Figure S13). The community structure of *Ca*Mg, on the other hand, was significantly different between the sites (PERMANOVA, *P*<0.01) and also clustered significantly by host genotype (PERMANOVA, *P*<0.01) (Figure 3). Indeed, close inspection of the PCoA plot indicated that the separation of *Ca*Mg communities by site is actually due to taxonomic affiliation of the fungal spores which themselves differed by location. This is best visible at the bottom of the plot where*Ca*Mg communities from VTU *Scutellospora* cluster together even though they came from different sampling sites. To better understand the relationship between AMF host identity and *Ca*Mg population diversity, we calculated various phylogenetic diversity metrics of *Ca*Mg populations found in individual AMF spores. We found that phylogenetic diversity was lower and less consistent in VTU *Acaulospora* when compared among the four most abundant AMF VTUs (Figure S13). Analogously, *Ca*Mg sequences were less dispersed (more similar) in VTUs *Acaulospora* and *Diversispora*. Together, these results indicate that AMF host genotype plays a role in structuring *Ca*Mg communities.

Only 7 spores in our dataset were exclusively populated by *Ca*Mg, whereas the majority (n=80) harboured at least one other endobacterial OTU. To understand if there was any relationship between *Ca*Mg and non-*Ca*Mg sequences within AMF spores, we plotted their respective diversity indices. Interestingly, we found a significant correlation between the diversity of *Ca*Mg and the diversity of non-*Ca*Mg OTUs within spores. As *Ca*Mg diversity increased, diversity of other endobacteria decreased (Figure 5). This pattern is indicative of competition dynamics between *Ca*Mg and non-*Ca*Mg endobacterial populations and might provide insight into the role of *Ca*Mg symbionts in AMF.

**Figure 5.**
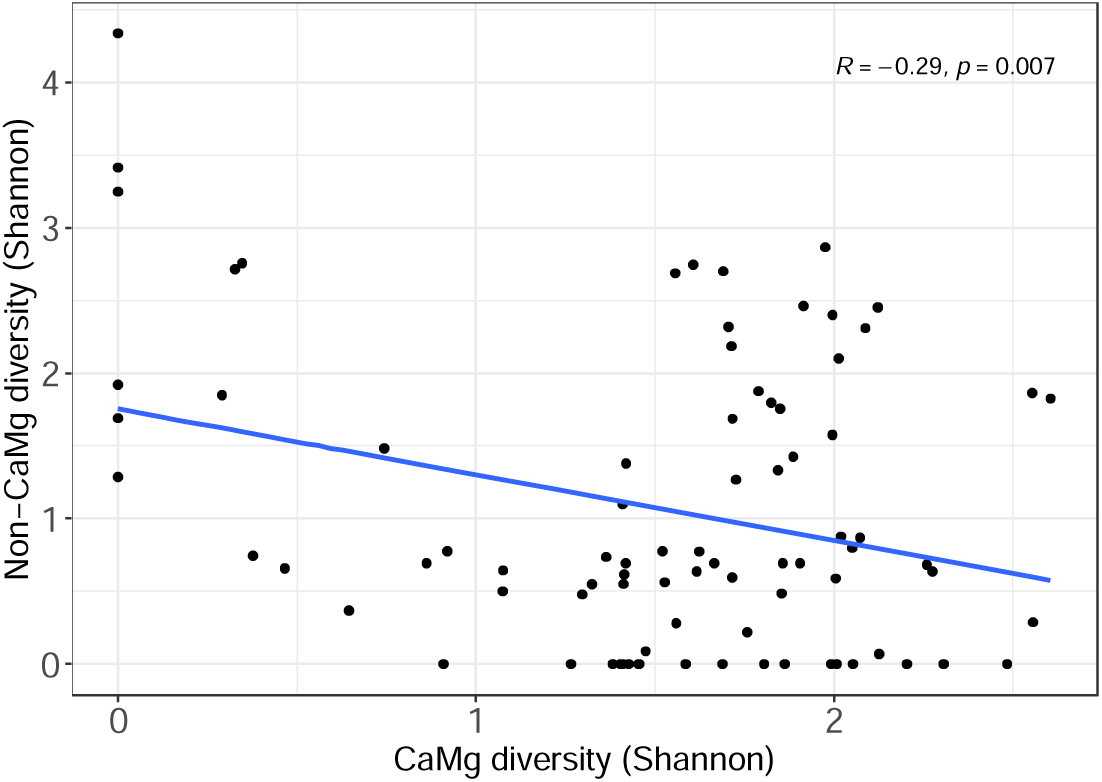
Significant correlation between *Ca*Mg and non-*Ca*Mg Shannon diversity. Pearson correlation coefficient and P value were calculated by the stat_cor function in R, grey area represents 95% confidence interval.

## Discussion

In this study we characterized the entire diversity of bacteria co-existing inside environmental spores of AMF and in doing so, identified hundreds of bacterial taxa, previously unknown to reside within AMF. Despite this unprecedented diversity, the most prevalent and abundant endobacteria were the known endosymbiotic *Ca*Mg, whose role in AMF biology remains experimentally untested. By analyzing the population structure of *Ca*Mg as well as the other co-occurring AMF endobacteria, we formulated a hypothesis centered around a conditional mutualistic relationship between *Ca*Mg and AMF.

Consistent with previous reports, *Ca*Mg populations were highly heterogenous within individual AMF spores, with as many as 50 OTUs detected in our study. Such population heterogeneity within individual hosts is considered a hallmark of parasites^15–17^ due to predicted increase in competition dynamics among symbionts which could be detrimental to their host. However, we hypothesize that this population heterogeneity could be an evolutionary adaptation for outcompeting other bacteria. The incidence of *Ca*Mg in field AMF spores is extremely high, ranging from 70-88% (^12^ and this study). Interestingly, this incidence does not differ across different ecosystems (agricultural grassland versus dunes) or between dune environments separated by thousands of miles (Irish vs American North Atlantic dunes^12^), indicating an unknown functional significance of these incidence rates. It seems very unlikely that *Ca*Mg would persist in AMF populations over evolutionary time to such high levels if they were not providing any benefit to AMF. In fact, according to evolutionary theory, the ability to act as *conditional mutualists* could account for the persistence of *Ca*Mg symbionts in AMF populations^47–49^. In a *conditional mutualism*, symbionts improve host fitness under specific conditions, contingent on the ecological context, symbiont’s life history stage and population size^50^. In this way, cost and benefit of harboring symbionts may fluctuate depending on the environment. For example, when symbionts provide protection to their host from predation, benefit of harboring the symbionts is high when predators are present but decreases when they are absent*i.e.* benefit is *conditional* to presence of predators. In the world of bacterial-fungal interactions it has recently come to light that fungi <u>do</u> in fact harbor bacterial endosymbionts for protection against predators. The fungus *Mortierella* harbours *Mycoavidus* endobacteria which produce toxins that protect their host from predation by nematodes^51^. Analogously, *Mycetohabitans* endobacteria protect their *Rhizopus microsporus* fungal host from predation by amoebae and nematodes^52^. Our discovery that spores with higher *Ca*Mg diversity harbour less diverse communities of other endobacteria provide a glimpse into how *Ca*Mg might benefit their AMF host. By maintaining high population diversity within AMF, *Ca*Mg are likely more effective at outcompeting other endobacteria which might otherwise be detrimental to AMF. In this way, by tolerating highly diverse *Ca*Mg populations, an AMF individual would still be home only to one group of endosymbionts belonging to the same genus. This would mean that any competition dynamics between closely related *Ca*Mg are only marginally costly as compared to competitive dynamics between endobacteria from different phyla. We therefore propose that*Ca*Mg are likely “conditional mutualists” of AMF, analogously to defensive mutualists of *R. microsporus* and *Mortierella*. Of course, this requires further experimental confirmation.

The discovery that *Ca*Mg incidence rate, intra-spore relative abundance (Figure S9), species richness (Figure S13), community structure (Figure 3) and phylogenetic diversity (Figure S13) are all related to AMF host identity, indicates that host genotype is a significant driver of *Ca*Mg endobacterial population assembly and that different *Ca*Mg populations are associated with different AMF taxa. This is analogous to what has previously been observed in plants^53^, animals^54, 55^ and more recently in fungal fruiting bodies^56^ whereby host genetics are a main driver of microbiome community structure. Given that the study of fungi-associated bacteria is a growing field of research^57, 58^ this work provides important insights into how fungal microbiomes are assembled.

Our findings that in addition to known AMF endosymbionts, many other phylogenetically diverse bacteria can be found inside AMF spores was surprising. Why would these bacteria, most of which are not known to have an endosymbiotic lifestyle, want to exist inside AMF spores? Recent research is suggesting that bacteria can exploit fungi for their shelter, facilitating survival under unfavorable conditions^59^. Diverse bacteria were shown to colonize thick-walled fungal resting structures known as chlamydospores and had higher survival rates than planktonic bacteria under abiotic stress. Interestingly, bacteria which entered chlamydospores were not known to have an endosymbiotic lifestyle and instead represented diverse species of free-living soil dwellers as well as plant and animal pathogens^59^. These new findings have profound ecological implications by providing an explanation of how free-living bacteria survive under unfavorable conditions such as low nutrient availability and periods of freezing^60^. In light of these findings, we speculate that the diverse bacterial communities detected in AMF spores as part of this study might also include these transient hitchhikers which exploit the lipid-rich AMF spores as a source of shelter.

Unlike *Ca*Mg, *Ca*Gg were extremely rare, consistent with previous reports that these endobacteria are not widely distributed in environmental AMF isolates^12^. Our PCR based screen detected *Ca*Gg in only 2 spores, whereas deep sequencing detected *Ca*Gg in 10 spores, highlighting the difference in sensitivity between these two methods and/or primer sets. Remarkably, we detected *Ca*Gg in spores of *Paraglomeracea* and *Ambisporaceae* (Table S2), whereas *Ca*Gg have previously only been reported in *Gigasporacea*^13^. This indicates that *Ca*Gg endosymbionts might have a wider host range than previously thought. Moreover, we detected a potentially novel lineage of Mollicutes-related bacteria inside AMF. These bacteria were distinct from all known Mollicutes and likely represent a previously unknown endosymbiont of AMF as well as a novel lineage of Mollicutes (Figure S12).

Lastly, this study analyzed the effect of different measured environmental parameters on *Ca*Mg distribution in AMF spores. In site 1, which was an agricultural grassland subject to different levels of long term P fertilization, soil nutrient levels, including available P, significantly affected *Ca*Mg distribution in AMF. On the other hand, at site 2 (dune) plant density was the main driver of *Ca*Mg incidence in AMF spores. Plant density (in the dunes) and soil P (in the agricultural grassland) were also important drivers of root endosphere AMF communities at these sites^22^ therefore it is possible that by influencing the structure of AMF communities, the effect of plant density and P translated into *Ca*Mg distribution across AMF spores at these sites.

Taken together, our study highlights a previously unknown complexity of AMF-associated endobacteria and emphasizes the need to understand their roles in the biology of these ubiquitous plant root symbionts. Future research should be directed at unravelling the functions of different endobacterial communities in AMF biology and how they might affect the interaction between AMF and their plant hosts. Furthermore, additional ecological studies are needed to understand the global distribution and diversity of AMF endobacteria, particularly focusing on different plant host species. Such research will further our understanding of AMF-plant-bacterial interactions and would be applicable in informing soil conservation efforts as well as the use of AMF in sustainable agriculture.

## Data availability

All sequencing data has been submitted to the Sequence Read Archive (SRA) National Center for Biotechnology Information (NCBI) database under the project number PRJNA986508.

## Acknowledgements

We thank Marc Barker for help with field sampling, Jim Grant for advice on statistical analysis, Teresa Pawlowska for comments on an earlier version of this manuscript and Graham Hughes for advise on phylogenetic analysis. We would also like to acknowledge John Murphy for maintaining the long term agricultural grassland site and the technical staff at Johnstown Castle Research laboratories for soil chemical analysis. This work was funded by the Government of Ireland Postdoctoral Fellowship Programme (Project no. GOIPD/2017/879).

## Author Contributions

O.A.L., F.P.B., D.W. and E.D. designed research; O.A.L., F.P.B. and D.W. carried out field work; O.A.L, and S.P. performed experiments; O.A.L. and T.C. analyzed data; O.A.L. and T.C. wrote the paper.

