## Supplementary material for "Community structure of known and previously unknown endobacteria associated with spores of arbuscular mycorrhizal fungi": Figure S1

**Supplementary Information**

Figure S1
Figure S2
Figure S3
Figure S4
Figure S5
Figure S6
Figure S7
Figure S8
Figure S9
Figure S10
Figure S11
Figure S12
Figure S13

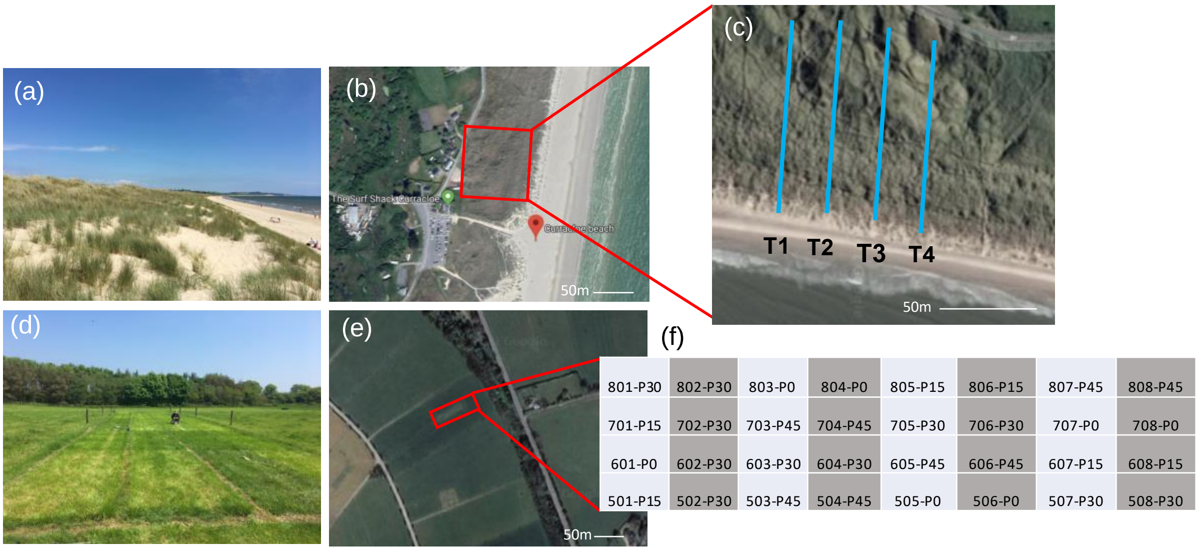


**Figure S1.** Sampling sites and sampling design. (a) Site 2 photo; (b) site 2 satellite view from Google Maps taken in 2020; (c) configuration of the site 2 sampling transects (blue lines) Transect 1-Transect 4 denoted T1-T4; (d) site 1 site photo; (e) site 1 satellite view from Google maps taken in 2020; (f) configuration of the site 1 sampling plots, light blue boxes denote "-slurry plots", grey boxes denote "+slurry" plots. Numbers correspond to plot number followed by P fertilization level from 0-45 kg $Ha^{-1}$ $Yr^{-1}$. Site 1, agricultural grassland; site 2, dune. Reproduced with permission from Lastovetsky *et al*. 2022 (https://doi.org/10.1111/1462-2920.16227).

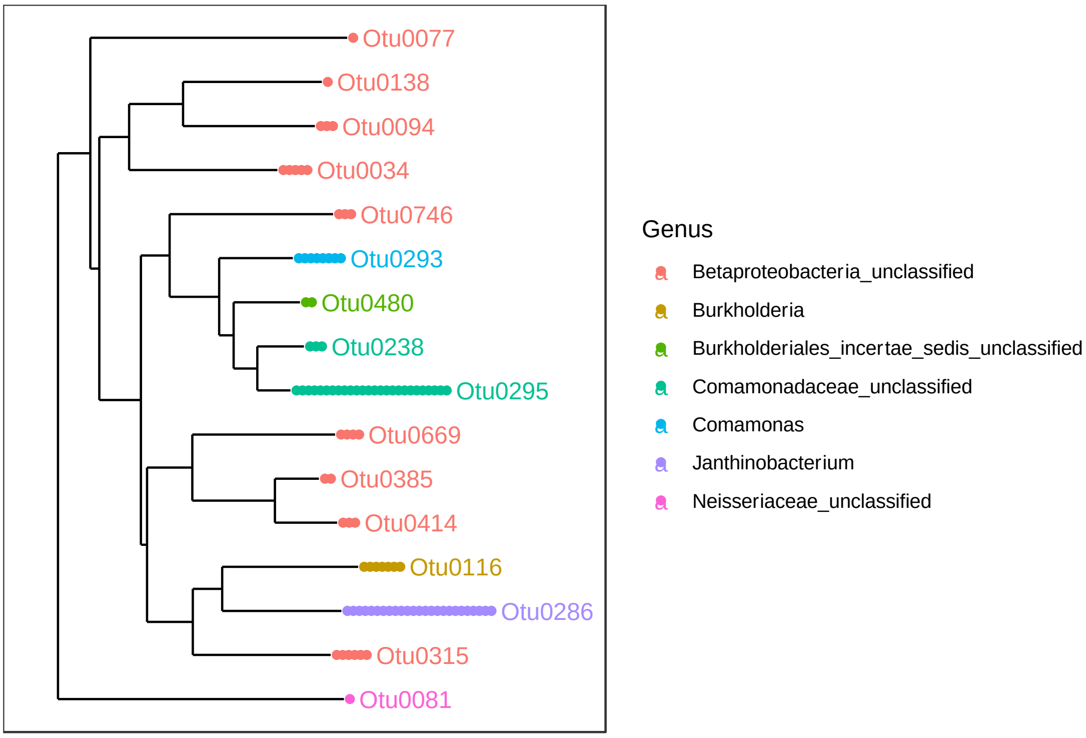


**Figure S2**. Bacterial OTUs belonging to "Betaproteobacteria" used to identify potential *Ca*Gg OTUs. Neighbour joining tree was created for OTUs with >30 reads through the phyloseq R package.

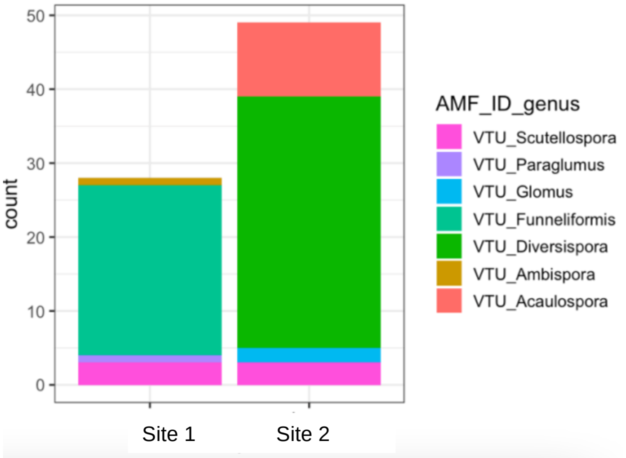


**Figure S3.** Relative abundance of AMF VTUs at the sampling sites. The bars are of uneven heights due to less spores being recovered from the agricultural grassland site.

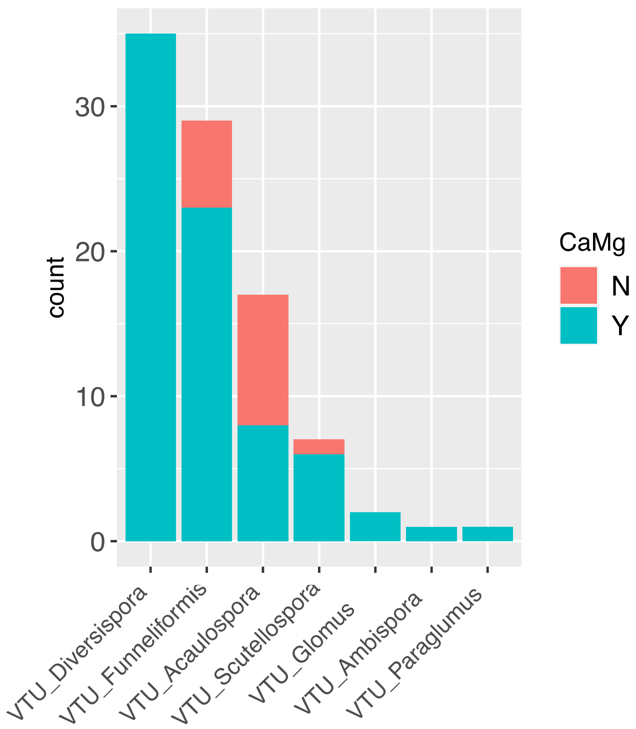


**Figure S4.** Incidence of *Ca*Mg endobacteria in the different AMF VTUs. N = not present, Y = present. Y-axis shows number of spores (i.e. count).

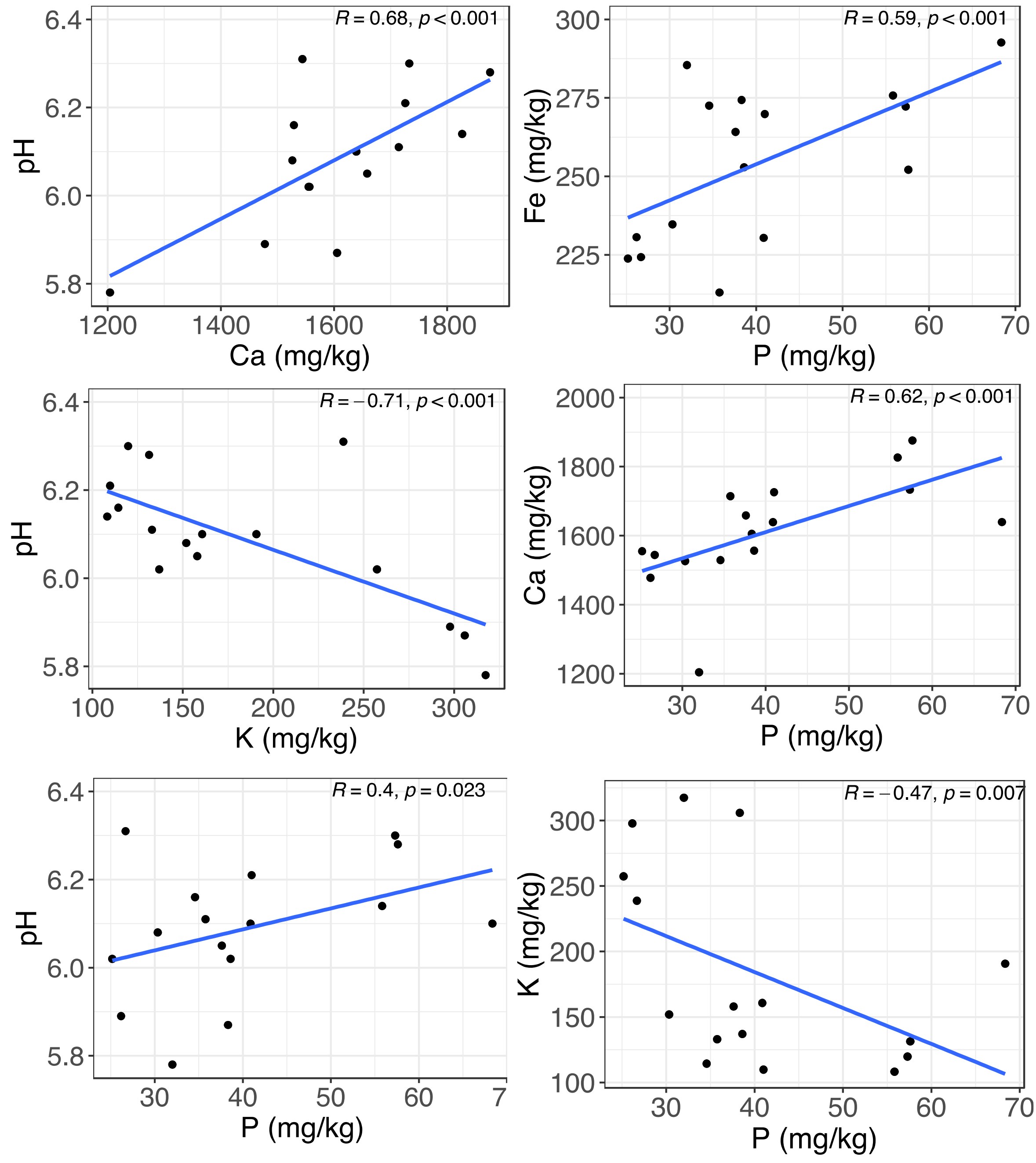


**Figure S5.** Significant correlations between soil parameters at site 2 (agricultural grassland). Pearson correlation coefficient and *P* value were calculated by the stat_cor function in R.

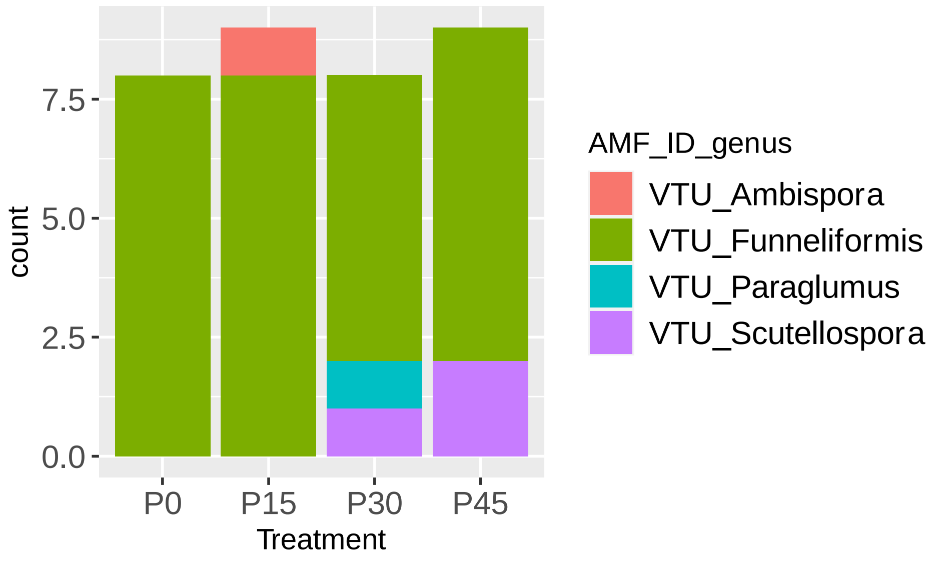


**Figure S6.** Distribution of AMF VTUs across sampling points subject to different P treatments. P0, P15, P30 and P45 received 0, 15, 30 and 45 kg $ha^{-1}$ $yr^{-1}$ respectively.

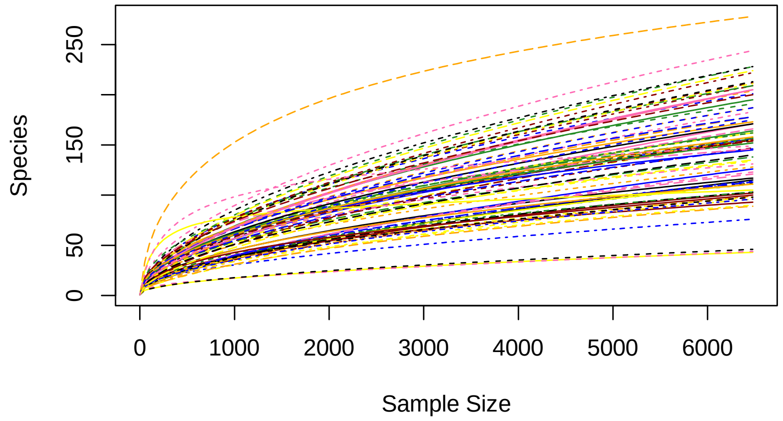


**Figure S7.** Bacterial OTU (based on 16S) accumulation curves per sample within AMF spores.

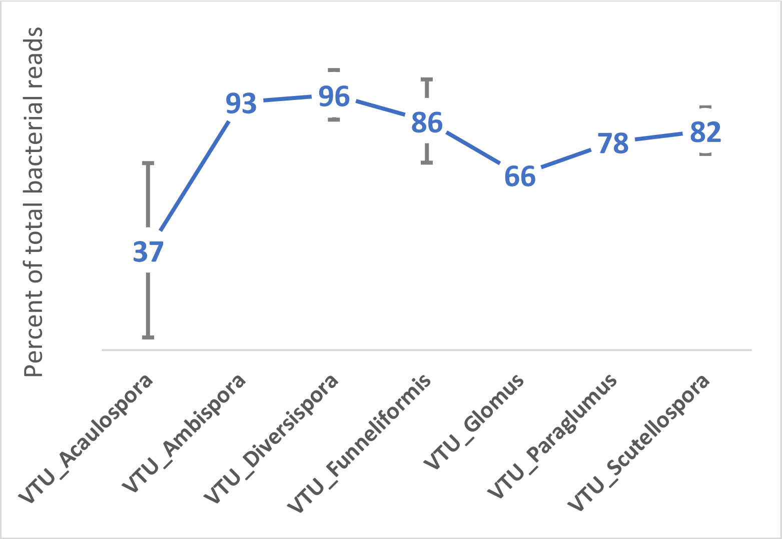


**Figure S8.** Percentage of reads belonging to *Ca*Mg endobacteria across different AMF VTUs. Error bars represent 1 SEM.

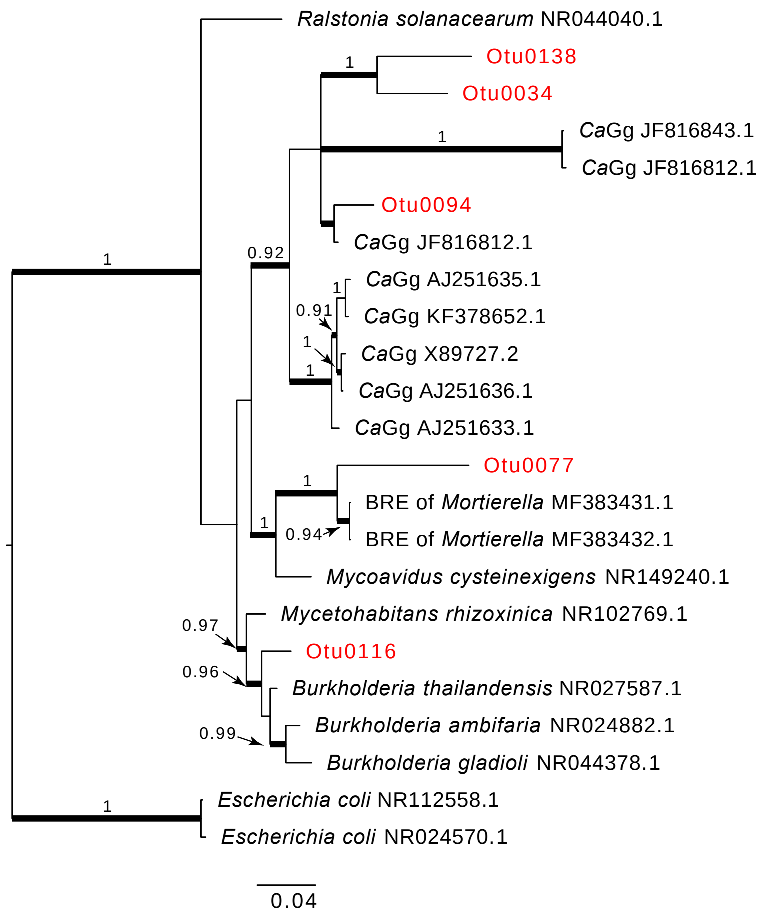


**Figure S9:** Relationships between *Ca*Gg, *Burkholderia*-related endobacteria of *Rhizopus microsporus* (*Mycetohabitans rhizoxinica*) and *Mortierella elongata* (*Burkholderia*-related endobacteria, BRE, and *Mycoavidus cysteinexigens*), as well as free-living *Burkholderia* based on the 16S rRNA gene sequence. Taxon identifiers in red represent bacterial sequences obtained from this study which had hits to known *Ca*Gg sequences in the nt BLAST database. Bayesian posterior probability values above 0.8 are displayed above branches, branches with Maximum Likelihood bootstrap support over 70% are thickened. The tree was rooted with *E. coli*.

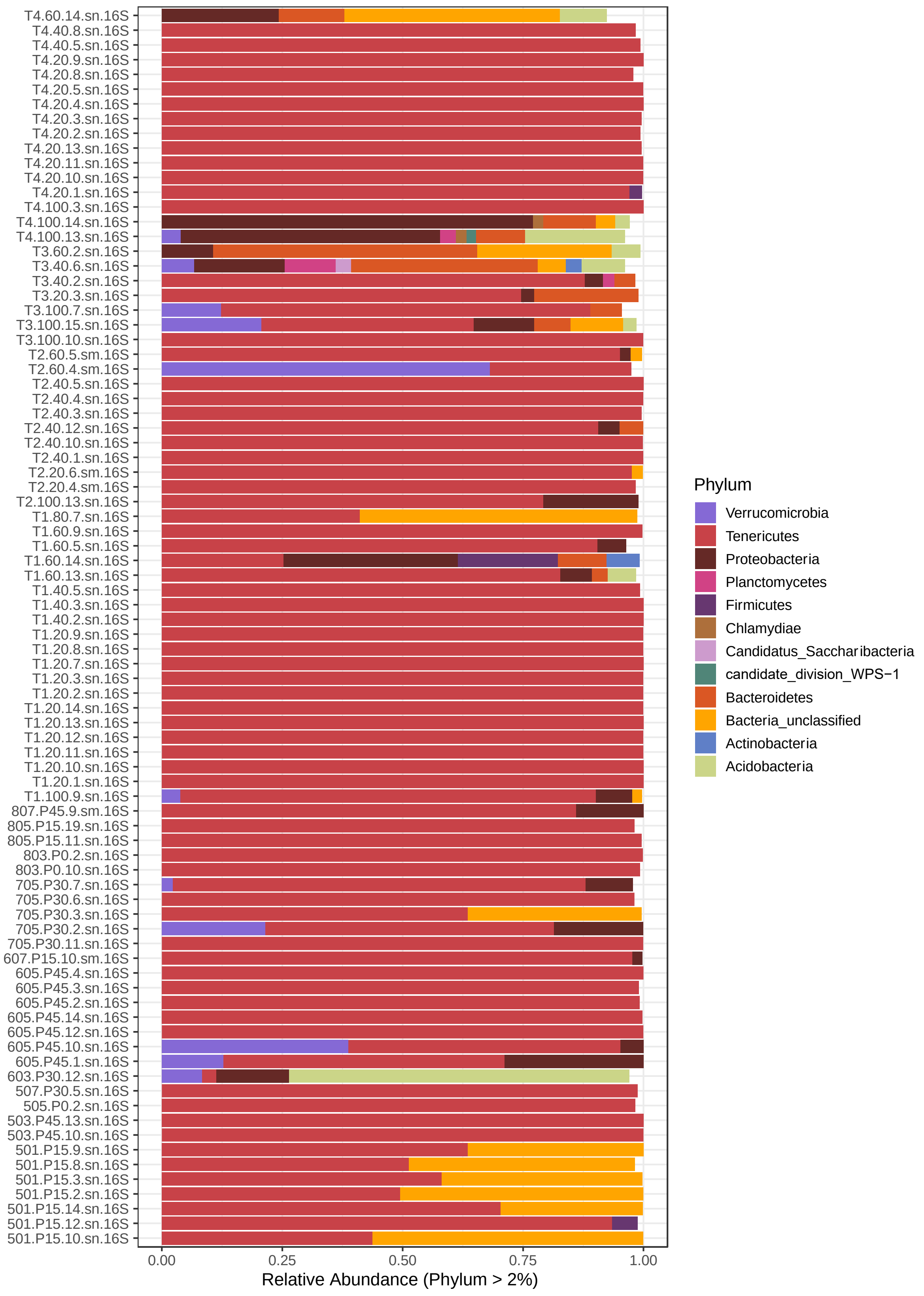


**Figure S10.** Relative Abundance of endobacteria at Phylum level in all AMF spore samples. Each sample name on the y-axis corresponds to an individual AMF spore. Sample names starting with a number are from site 1 (agricultural grassland); sample names starting with a "T" are from site 2 (dune); "sn" in the sample name corresponds to "spore isolated in November"; "sm" corresponds to "spore isolated in March".

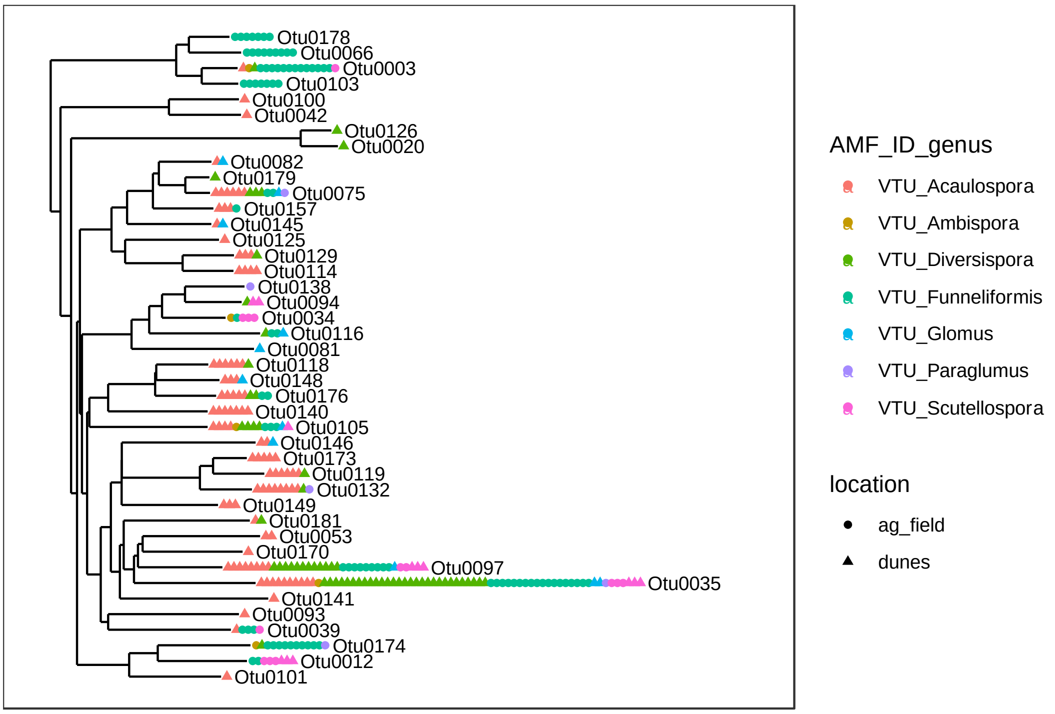


**Figure S11**. Distribution of the non-*Ca*Mg bacterial OTUs across AMF VTUs and sampling sites. Neighbour joining tree was created for OTUs with >500 reads thought the phyloseq R package.

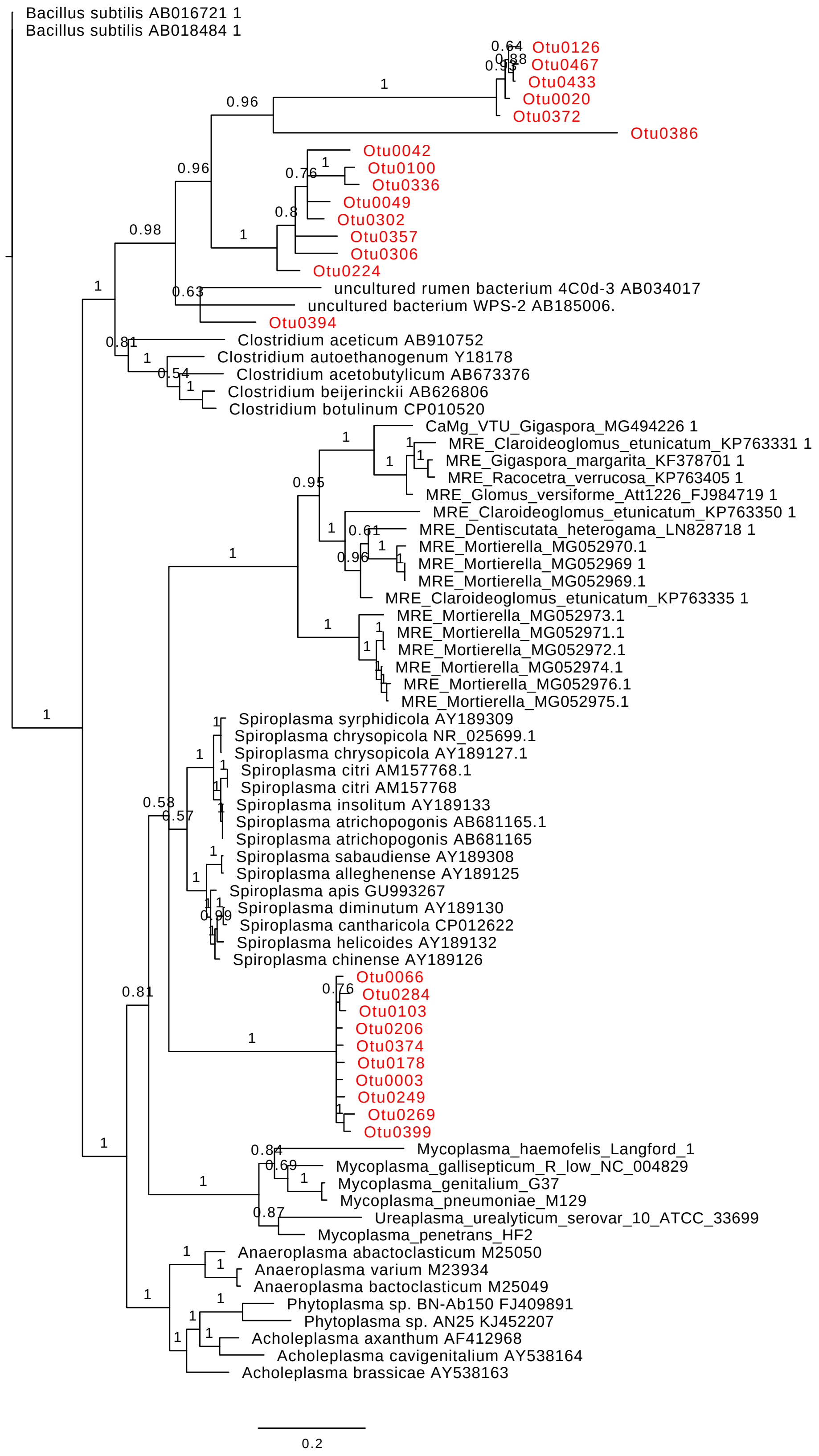


**Figure S12.** Phylogenetic placement of 'unclassified" bacterial OTUs. Taxa labelled in red represent the unclassified bacterial OTUs from this study. Bayesian phylogeny reconstruction was performed based on 16S rRNA gene sequence. Posterior probability values are displayed above branches.

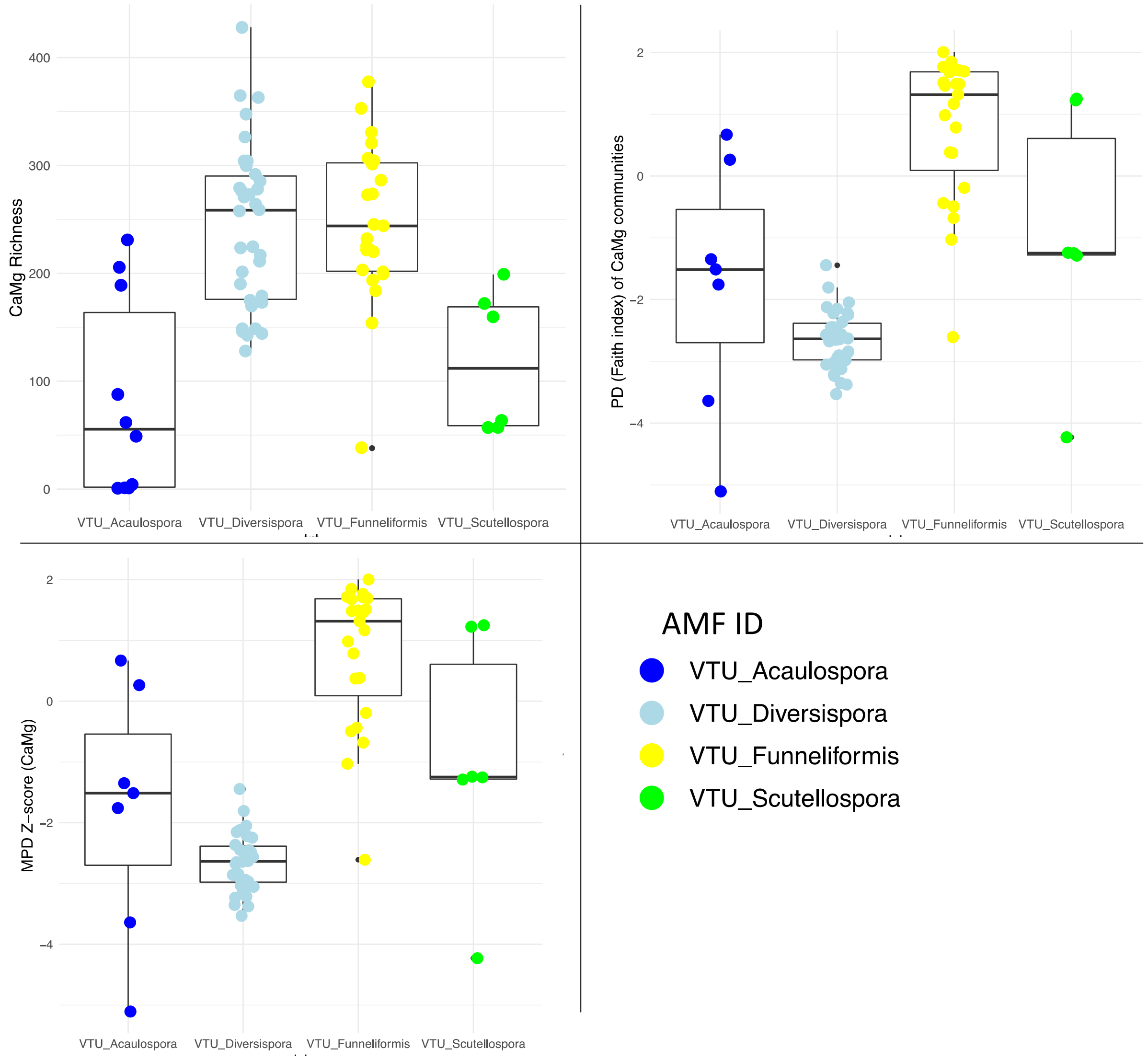


**Figure S13.** Diversity of *Ca*Mg populations in the top 4 most abundant AMF VTUs. *Top left pane: Ca*Mg richness (OTU count); *top right pane: Ca*Mg phylogenetic diversity (PD, *i.e.* Faith Index); *bottom left pane: Ca*Mg standardized effect size mean phylogenetic (pairwise) distances (MPD z score). MPD_z indicates measures clustering or dispersion of communities, with more negative values indicating clustering.
