## Supplementary material for "Community structure of known and previously unknown endobacteria associated with spores of arbuscular mycorrhizal fungi": Table S1

**Table S1. Soil pH and nutrient levels at the dune and agricultural grassland sampling points.** P treatment= Phosphorus fertilizer levels applied in kg Ha^-1^ Yr^-1^. Nutrient levels (P, S, K, Mg, Al, Fe, Ca, Co, Ci, Mn, Zn) in mg kg^-1^. This table is adapted from Lastovetsky et al. 2022 https://doi.org/10.1111/1462-2920.16227.

| Location | Sample | P Treatment | pH | P | S | K | Mg | Al | Fe | Ca | Co | Cu | Mn | Zn |
| --- | --- | --- | --- | --- | --- | --- | --- | --- | --- | --- | --- | --- | --- | --- |
| Dune (site 2) | T1 20 | 0 | 8.89 | 2.4 | 2.25 | 9.87 | 34.81 | <5 | 18.12 | 1245.21 | <0.1 | 0.1 | 10.67 | <1 |
| Dune (site 2) | T1 40 | 0 | 8.42 | 1.61 | 4.23 | 9.13 | 39.66 | <5 | 14.88 | 1475.44 | <0.1 | <0.1 | 9.45 | <1 |
| Dune (site 2) | T1 60 | 0 | 8.5 | 1.84 | 2.39 | 9.98 | 34.8 | <5 | 19.06 | 1367.76 | <0.1 | 0.11 | 11.08 | <1 |
| Dune (site 2) | T1 80 | 0 | 8.23 | 1.61 | 3.9 | 7.31 | 41.42 | <5 | 20.79 | 1721.43 | <0.1 | 0.11 | 11.46 | 1.1 |
| Dune (site 2) | T1 100 | 0 | 8.34 | 2.16 | 3.44 | 16.46 | 37.65 | <5 | 28.48 | 1325.01 | <0.1 | 0.15 | 14.33 | 2.18 |
| Dune (site 2) | T2 20 | 0 | 8.99 | 1.97 | 4.58 | 11.24 | 37.3 | <5 | 16.96 | 1138.69 | <0.1 | 0.05 | 9 | <1 |
| Dune (site 2) | T2 40 | 0 | 8.95 | 1.13 | 2.5 | 8.69 | 28.11 | <5 | 14.14 | 1272.95 | <0.1 | 0.09 | 9 | <1 |
| Dune (site 2) | T2 60 | 0 | 8.55 | 1.34 | 4.1 | 2.82 | 31.87 | <5 | 16.37 | 1654.36 | <0.1 | 0.06 | 10.1 | <1 |
| Dune (site 2) | T2 80 | 0 | 8.4 | 1.65 | 0.47 | 14.96 | 45.66 | <5 | 22.91 | 1592.69 | <0.1 | 0.14 | 12.02 | 1 |
| Dune (site 2) | T2 100 | 0 | 8.35 | 0.72 | 3.3 | 4.09 | 48.4 | <5 | 21.11 | 2940.63 | <0.1 | <0.1 | 10.72 | <1 |
| Dune (site 2) | T3 20 | 0 | 8.72 | 2.72 | 3.19 | 17.01 | 34.36 | <5 | 14.98 | 919.87 | <0.1 | <0.1 | 9.35 | <1 |
| Dune (site 2) | T3 40 | 0 | 8.68 | 1.2 | 3.17 | 2.45 | 24.85 | <5 | 15.5 | 1369.34 | <0.1 | <0.1 | 9.6 | <1 |
| Dune (site 2) | T3 60 | 0 | 8.5 | 1.47 | 2.44 | 13.71 | 43.28 | <5 | 18.68 | 2148.09 | <0.1 | 0.11 | 10.21 | <1 |
| Dune (site 2) | T3 80 | 0 | 8.31 | 0.83 | 5.04 | 5.1 | 37.97 | <5 | 21.66 | 2091.15 | <0.1 | 0.08 | 11.22 | <1 |
| Dune (site 2) | T3 100 | 0 | 8.06 | 3.35 | 2.71 | 7.78 | 35 | <5 | 28.78 | 634.27 | <0.1 | 0.13 | 12.94 | 2.51 |
| Dune (site 2) | T4 20 | 0 | 8.84 | 1.93 | 4.55 | 6.04 | 31.26 | <5 | 15.7 | 967.75 | <0.1 | 0.05 | 9.79 | <1 |
| Dune (site 2) | T4 40 | 0 | 8.76 | 1.13 | 5.01 | 5.34 | 26.94 | <5 | 14.38 | 1423.2 | <0.1 | 0.04 | 9.43 | <1 |
| Dune (site 2) | T4 60 | 0 | 8.69 | 2.15 | 3.1 | 5.58 | 31.43 | <5 | 17.31 | 1956.66 | <0.1 | <0.1 | 11.4 | <1 |
| Dune (site 2) | T4 80 | 0 | 8.6 | 1.16 | 2.91 | 16.3 | 28.92 | <5 | 19.16 | 1206.31 | <0.1 | 0.11 | 10.86 | <1 |
| Dune (site 2) | T4 100 | 0 | 8.18 | 3.46 | 4.46 | 6.11 | 41 | <5 | 23.1 | 485.32 | <0.1 | 0.11 | 11.25 | 1.69 |
| Ag (site 1) | 501 | 15 | 5.87 | 38.32 | 23.32 | 305.85 | 63.71 | 566.25 | 274.33 | 1605.43 | 0.13 | 2.05 | 14.6 | 2.91 |
| Ag (site 1) | 601 | 0 | 5.78 | 32 | 32.15 | 317.42 | 56.13 | 567.5 | 285.45 | 1204.12 | 0.12 | 1.47 | 12.65 | 1.58 |
| Ag (site 1) | 701 | 15 | 6.16 | 34.57 | 21.7 | 114.37 | 57.14 | 545.26 | 272.53 | 1529.05 | 0.14 | 1.39 | 14.56 | 1.13 |
| Ag (site 1) | 801 | 30 | 6.21 | 41 | 30.36 | 109.81 | 59.96 | 580.95 | 269.85 | 1725.85 | 0.13 | 1.32 | 13.74 | 1.36 |
| Ag (site 1) | 503 | 45 | 6.14 | 55.83 | 23.42 | 108.22 | 73.83 | 566.37 | 275.76 | 1826.39 | 0.15 | 2.11 | 19.1 | 1.75 |
| Ag (site 1) | 603 | 30 | 6.02 | 38.61 | 19.15 | 136.98 | 61.64 | 545.4 | 252.9 | 1556.54 | 0.2 | 1.59 | 22.46 | 1.6 |
| Ag (site 1) | 703 | 45 | 6.3 | 57.3 | 19.37 | 119.76 | 58.14 | 546.38 | 272.24 | 1733.02 | 0.2 | 1.5 | 23.59 | 1.49 |
| Ag (site 1) | 803 | 0 | 6.02 | 25.17 | 23.15 | 257.36 | 73.54 | 593.44 | 223.81 | 1554.91 | 0.23 | 1.51 | 29.33 | 2.45 |
| Ag (site 1) | 505 | 0 | 5.89 | 26.17 | 31.39 | 297.76 | 67.58 | 551.11 | 230.61 | 1477.78 | 0.23 | 1.7 | 38.71 | 1.85 |
| Ag (site 1) | 605 | 45 | 6.1 | 68.35 | 19.09 | 190.69 | 60.81 | 564.28 | 292.64 | 1639.4 | 0.21 | 1.55 | 36.94 | 1.68 |
| Ag (site 1) | 705 | 30 | 6.1 | 40.87 | 22.23 | 160.7 | 62.95 | 569.26 | 230.41 | 1639.05 | 0.3 | 1.54 | 47.6 | 2.55 |
| Ag (site 1) | 805 | 15 | 6.11 | 35.76 | 33.07 | 132.99 | 64.57 | 579.36 | 212.98 | 1714.35 | 0.3 | 1.53 | 45.71 | 3.85 |
| Ag (site 1) | 507 | 30 | 6.05 | 37.62 | 26.69 | 157.98 | 63.27 | 583.46 | 264.16 | 1658.55 | 0.37 | 1.89 | 66.95 | 1.64 |
| Ag (site 1) | 607 | 15 | 6.08 | 30.33 | 26.95 | 151.91 | 52.29 | 568.51 | 234.7 | 1526.35 | 0.41 | 1.5 | 74.29 | 1.74 |
| Ag (site 1) | 707 | 0 | 6.31 | 26.68 | 21.09 | 238.77 | 50.06 | 543.17 | 224.28 | 1544.15 | 0.43 | 1.36 | 80.95 | 2.86 |
| Ag (site 1) | 807 | 45 | 6.28 | 57.61 | 30.25 | 131.36 | 59.93 | 595.67 | 252.11 | 1875.78 | 0.32 | 1.21 | 55.95 | 1.56 |
